# Construction of a high-density American cranberry (*Vaccinium macrocarpon* Ait.) composite map using genotyping-by-sequencing for multi-pedigree linkage mapping

**DOI:** 10.1101/088419

**Authors:** Brandon Schlautman, Giovanny Covarrubias-Pazaran, Luis Diaz-Garcia, Massmo Iorizzo, James Polashock, Edward Grygleski, Nicholi Vorsa, Juan Zalapa

## Abstract

The American cranberry (*Vaccinium macrocarpon* Ait.) is a recently domesticated, economically important, fruit crop with limited molecular resources. New genetic resources could accelerate genetic gain in cranberry through characterization of its genomic structure and by enabling molecular-assisted breeding strategies. To increase the availability of cranberry genomic resources, genotyping-by-sequencing (GBS) was used to discover and genotype thousands of single nucleotide polymorphisms (SNPs) within three inter-related cranberry full-sib populations. Additional simple sequence repeat (SSR) loci were added to the SNP datasets and used to construct bin maps for the parents of the populations, which were then merged to create the first high-density cranberry composite map containing 6073 markers (5437 SNPs and 636 SSRs) on 12 linkage groups (LGs) spanning 1124 cM. Interestingly, higher rates of recombination were observed in maternal than paternal gametes. The large number of markers in common (mean of 57.3) and the high degree of observed collinearity (mean Pair-wise Spearman Rank Correlations > 0.99) between the LGs of the parental maps demonstrates the utility of GBS in cranberry for identifying polymorphic SNP loci that are transferable between pedigrees and populations in future trait-association studies. Furthermore, the high-density of markers anchored within the component maps allowed identification of segregation distortion regions, placement of centromeres on each of the 12 LGs, and anchoring of genomic scaffolds. Collectively, the results represent an important contribution to the current understanding of cranberry genomic structure and to the availability of molecular tools for future genetic research and breeding efforts in cranberry.

## INTRODUCTION

The American cranberry (*Vaccinium macrocarpon* Ait.) is a diploid woody perennial in the Ericaceae family, and like other Ericaceous species, it grows well in acidic, nutrient poor soils (Vander Kloet 1983b; Kron *et al.* 2002; Vander Kloet and Avery 2010). Within the Ericaceae family, the cranberry and blueberry species of sections *Oxycoccus* (Hill) Koch and *Cyanococcus* A. Gray share a common base chromosome number (n=12), and a similar karyotype with relatively uniformly sized metacentric chromosomes (Camp 1945; Hall and Galleta 1971; Vander Kloet 1983a, 1983b; Costich *et al.* 1993). Both crops are growing in popularity and commercial importance because of the numerous health benefits attributed to their phytochemical constituents, and global production of cranberry and blueberry has expanded to encompass a combined 100,000+ hectares valued at more than 1.5 billion U.S. dollars (Howell *et al.* 1998; Foo *et al.* 2000; Duarte *et al.* 2006; FAO 2012; Feghali *et al.* 2012). However, cranberry genetic improvement has lagged behind its blueberry brethren for various reasons including: the lack of continued breeding efforts, the limited number of breeding programs in existence, the long breeding cycle due to lengthy establishment periods (2-4 years) followed by long evaluation periods (4-5 years) required to measure biennial bearing (Eaton *et al.* 1983; Elle 1996; Vorsa and Johnson-cicalese 2012; Schlautman *et al.* 2015a), and a lack of adoption of molecular-assisted breeding strategies which could increase the rate of domestication and genetic gain in cranberry.

Prerequisites to marker-assisted breeding strategies, such as molecular-assisted seedling selection and/or genomic prediction (Ru *et al.* 2015; Covarrubias-Pazaran 2016), include the availability of numerous molecular markers distributed throughout the genome for haplotype estimation, knowledge of associations between markers and traits of interest, available populations with characterized genetic and phenotypic diversity, and high-throughput marker genotyping methodologies that justify the costs associated with molecular-assisted breeding compared to classical breeding methods. Large-scale exploration and development of genetic and genomic resources in cranberry has only begun recently with the advent of next generation sequencing technologies (NGS). For example, NGS technologies such as pyrosequencing (i.e., 454), sequencing by oligonucleotide ligation and detection (SOLiD), and Illumina have been useful in assembling both the plastid and mitochondrial genomes (Fajardo *et al.* 2013, 2014), a transcriptome (Polashock *et al.* 2014), and a draft nuclear genome assembly (Polashock *et al.* 2014). Simple sequence repeat (SSR) mining of those sequence resources resulted in the development of more than 900 informative (polymorphic) SSR markers (Georgi *et al.* 2011, 2013; Zhu *et al.* 2012; Schlautman *et al.* 2015b; Schlautman *et al.* 2016), with 136 and 541 of those SSRs placed in cranberry SSR linkage maps by (Georgi *et al.* 2013; Schlautman *et al.* 2015b), respectively. However, means for gathering SSR genotype information in an efficient, cost-effective manner at a breeding program scale have not been achieved in cranberry, and uncertainty remains about whether the available set of cranberry SSRs is large enough to saturate the genome.

Covarrubias-Pazaran *et al.* (2016) recently demonstrated the potential for genotyping-by sequencing in cranberry to serve as a high-throughput platform that integrates single nucleotide polymorphism (SNP) marker discovery and genotyping into a single procedure (Elshire *et al.* 2011). The Covarrubias-Pazaran *et al.* (2016) study increased the availability of cranberry DNA markers 10-fold and used them to construct the first high-density SNP cranberry linkage map. Therefore, to further saturate the cranberry genome with SNP markers and to test the utility of genotyping-by-sequencing to detect SNPs that are polymorphic across multiple populations, an experiment was designed to develop a cranberry composite map using genotyping-by-sequencing for multi-pedigree linkage mapping. In this study, six parental bin maps were constructed from three inter-related cranberry populations, whose ancestry trace back to seven historically important cranberry wild selections. The high-density of SNPs identified herein were placed in a composite map that allowed characterization of important chromosome structural aspects including identification of segregation distortion regions, centromere placement, and anchoring of cranberry nuclear scaffolds containing predicted coding DNA sequences.

## MATERIALS AND METHODS

### Plant Material and DNA extraction

Three full-sib linkage mapping populations (i.e. CNJ02, CNJ04, and GRYG) were derived from crosses between five inter-related cranberry parental genotypes (Figure 1). The CNJ02 population included 168 progeny from a cross between maternal parent, CNJ97-105 (*Mullica Queen^®^*), and paternal parent, NJS98-23 (*Crimson Queen^®^*); the CNJ04 population included 67 progeny from a reciprocal cross between CNJ97-105 (*Mullica Queen^®^*) and Stevens; and the GRYG population included 352 progeny from a cross between the maternal parent, [BGx(BLxNL)]95, and the paternal parent, GH1x35 (Figure 1). The CNJ02 and CNJ04 populations were generated and are maintained at the Rutgers University P.E. Marucci Center in Chatsworth, NJ and were planted in separate unreplicated, completely randomized designs. The GRYG population was generated and is maintained by the Valley Corporation in Tomah, WI. Genomic DNA from all parents and progeny was extracted from flash frozen newly emerged leaves using a Macherey-Nagel (MN) Plant II kit (Düren, Germany) following the manufacturer’s instructions.

**Figure 1.**
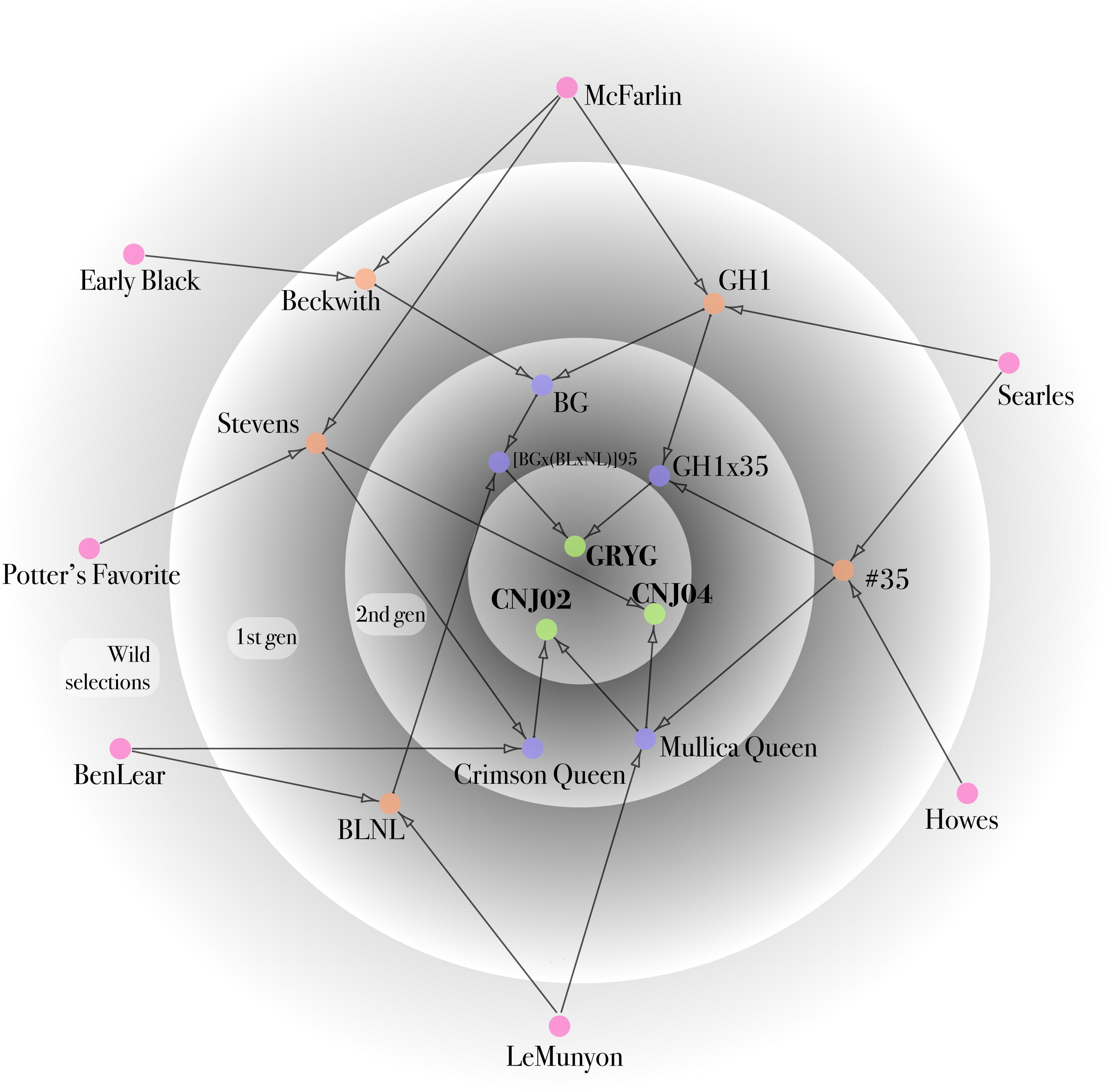
Description of the pedigrees of the three mapping populations (green), CNJ02, CNJ04, and GRYG, derived from crosses between five inter-related cranberry parental genotypes (blue) CNJ97-105 (*Mullica Queen^®^*), NJS98-23 (*Crimson Queen^®^*), Stevens, [BGx(BLxNL)]95, and GH1x35). All pedigrees trace to “The Big Seven” native cranberry selections (red), which have played important roles in the cranberry production and breeding history. The pedigree contains five first generation removed from the wild genotypes resulting from crosses between two native selections (orange).

### SSR and SNP genotyping and data processing

SSR marker data generated for the CNJ02 (541 SSRs) and GRYG (189 SSRs) populations in previous linkage mapping studies were incorporated into the current study (Schlautman *et al.* 2015a; Covarrubias-Pazaran *et al.* 2016). A subset of these previously mapped and additional unmapped SSR markers were used to genotype the CNJ04 progeny and its parents in this study (Schlautman *et al.* 2015a; Schlautman *et al.* 2015b). In addition, SSRs previously developed and mapped in blueberry (Rowland *et al.* 2014) and newly developed SSR primer pairs were used to genotype the CNJ02 progeny. Multiplex (3x) Polymerase Chain Reactions (PCR) and fragment analysis for SSR genotyping were performed according to (Schlautman *et al.* 2015a).

*Eco*T221-associated DNA fragments for the progeny and parents of the CNJ02 and CNJ04 mapping populations were generated and sequenced according to the genotyping-by-sequencing (GBS) approach described by Elshire *et al.* (2011). During library construction, adapters containing unique barcodes were ligated to restriction-digested DNA for the 235 progeny from the CNJ02 and CNJ04 populations and their 3 parental genotypes and then were pooled into three 96-plex sequencing libraries. To guarantee higher sequencing coverage in the parental genotypes, the parents for the CNJ02 and CNJ04 populations were represented by six samples each, and the resulting libraries were sequenced (single-end) on the Illumina HiSeq^®^ 2000 platform in the Cornell University Biotechnology Resource Center (BRC) Genomics Facility. Library preparation and sequencing for the GRYG population (352 progeny) was performed previously following the same methodology described herein, except the two parental genotypes were represented by four samples each. This sequence data, used in a previous publication (Covarrubias-Pazaran *et al.* 2016), was incorporated into the current study to increase the density of the final composite map and reprocessed to ensure that the linkage maps for all three populations were constructed according to the same standard set of parameters.

The reference-based Tassel v3.0.166 GBS analysis pipeline was used to filter and process the resulting sequence reads, align and merge sequence tags by genotype, and to call SNPs in the resulting datasets for the CNJ02 and CNJ04 populations using the parameters outlined in Glaubitz *et al.* (2014) (File S1). The reference genome used during SNP discovery and genotyping with the Tassel pipeline was created by concatenating the 229,745 scaffold cranberry genome assembly (N50=4,237 bp) produced by Polashock *et al.* (2014). After generating the filtered HapMap, the SNP marker data was further processed by removing SNPs separately from each of the three populations which had greater than 20% missing data, had a minor allele frequency (MAF) of less than 10 %, or had severely distorted segregation ratios.

### Parental Component Map Construction

Linkage analysis was performed with JoinMap v4.1 using the pseudo-testcross method. Markers were formatted and separated into two categories: a) uniparental markers heterozygous in only a single parent (lm x ll and nn x np); and b) biparental markers heterozygous in both parents (hk x hk, ef x eg, and ab x cd) (Van Ooijen 2006, 2011). Linkage groups (LGs) were determined with a LOD threshold > 5.0 (Van Ooijen 2006), and marker order was determined using the maximum likelihood (ML) algorithm (Van Ooijen 2011). A strict approach was used to determine and remove markers causing errors in estimation of marker ordering or inflation of map distances. First, markers which obviously caused problems and/or were placed far from the ends of the linkage groups (LGs) were identified visually when observing the phased genotype data. Additionally, iterative rounds of mapping were performed to remove markers that caused a nearest neighbor fit (cM) greater than 2 cM, a nearest neighbor stress greater than 0.035, or a nearest neighbor stress (cM) greater than 3.5 cM. Genetic distances among loci were then recalculated with the regression approach and the Kosambi mapping function using the fixed marker order determined by ML to facilitate map comparisons between the current maps and previous published maps.

Linkage mapping proceeded by first constructing a maternal and paternal component map for each population using only the uniparental markers (i.e. lm x ll or nn x np) and removing all markers causing problems in map order and distance per the guidelines outlined above. Using the resulting marker positional information, linkage-informed imputation was performed using the *linkim* package in R to impute missing marker information in the parental datasets (R Core Team 2015; Xu and Wu 2015). The multiple spanning tree (MST) algorithm implemented in the *ASMap* R package was then used to detect remaining genotyping errors and to perform bin mapping (determining bins of identical markers) for each of the parents (Wu *et al.* 2008; Taylor and Butler 2015).

Next, biparental markers were added to each parental map. For each population, a single uniparental marker from each bin of each parental bin map was added to the remaining biparental maker set (i.e hk x hk, ef x eg, and ab x cd markers) and linkage mapping was performed to construct integrated maps for the three populations using the before-mentioned guidelines with the *cp* approach in JoinMap v4.1. The resulting marker positional and phase information was used to convert the hk x hk biparental SNP markers to ab x cd maker format by linkage-informed imputation of the hk genotypes using functions from the *sommer* R package (Covarrubias-Pazaran 2016). Each biparental marker was then split into two uniparental markers (either lm x ll for alleles from the maternal parent or nn x np for alleles from the paternal parent) to create a new dataset for each parent containing all possible markers. The MST algorithm was used once again to detect genotyping errors within the biparental marker sets and to create bins for each of the parental maps using the Kosambi mapping function to estimate map distances (Wu *et al.* 2008; Taylor and Butler 2015). Pair-wise Spearman rank correlations comparing marker order in the LGs of the six parental component bin maps were estimated to ensure that they were syntenic, collinear and could be used in composite map construction. Additionally, collinearity between LGs was visually assessed in Circos circular ideograms generated in the R package *SOFIA* by plotting the links between homologous markers in the LGs of the parental bin maps in each of the three populations (Krzywinski *et al.* 2009).

### Composite Map Construction and Map Comparisons

A synthetic composite map was constructed from the six parental maps with the *LPmerge* package in R, which uses linear programming to minimize the mean absolute error between marker intervals in the parental maps and the composite map (Endelman and Plomion 2014). During composite mapping, iterations of the maximum interval size (*k*) ranging from *k* = 1 to *k* =10 were tested for each linkage group (LG), and the *k* which minimized the root mean square error (RMSE) when comparing the position of makers in the composite LG to the 6 parental LGs was chosen for construction of the final composite LG. Spearman rank correlations and visual assessment in Circos were used to determine the collinearity of the composite map with each of the 6 component bin maps, with the previous SSR map constructed map for the CNJ02 population (Schlautman *et al.* 2015a), and with the first GBS-based SNP map developed previously for the GRYG population (Covarrubias-Pazaran *et al.* 2016). Cranberry scaffolds from Polashock *et al.* (2014) containing predicted coding DNA sequences (CDS) and SSRs or SNPs mapped in this study were anchored using the markers’ positions in the composite map.

### Genome-wide segregation distortion and centromere placement

Segregation distortion for each mapped marker in parental bin maps was analyzed using chi-square tests (χ^2^) with one degree of freedom for codominant markers as implemented in JoinMap v4.1. The χ^2^ *p*-value for each locus in each parental component map was then plotted to examine patterns of segregation distortion and to determine if segregation distortion regions existed in any LGs. Markers with a χ^2^ *p*-value < 0.1 were considered distorted.

Centromeric regions in the cranberry linkage groups (LGs) of the component bin maps were explored and identified following the methodology of Limborg *et al.* (2016). Recombination frequencies (RF_M_) were estimated from phased genotype data by recording the proportion of offspring with an observed recombination (i.e. change of phase) in each interval between the terminal marker (m_0_) and every subsequent marker (m_n_) in both directions along the 12 cranberry LGs. For metacentric LGs, centromeric regions in the LGs were defined as the region from the point of intersection between the RF_M_ estimates made from each terminal marker extending outwards until reaching the first marker interval with an RF_M_ = 0.45 in both directions (Limborg *et al.* 2016).

### Data availability

File S2 contains marker positions in the composite linkage map and the six parental component bin maps. File S3 contains all SNP genotype data for the six parental component bin maps.

## RESULTS

### SSR and SNP Genotyping

Differing sizes of SSR marker datasets were available for mapping in each of the three populations. The CNJ02 population, which was used in the first high density SSR linkage mapping study had the largest amount of SSR data (629 SSRs) available for linkage mapping (Table 1; Table S1) (Schlautman *et al.* 2015a). The GRYG population, which was used in the first cranberry SNP genotyping study (Covarrubias-Pazaran *et al.* 2016), had a similar amount of SSR data compared to the CNJ04 population with 189 and 221 SSR markers, respectively (Table 1; Table S1). The CNJ04 population has never been used in a peer-reviewed study, and all SSR and SNP marker data for that population was generated herein using multiplex (3x) PCR reactions.

**Table 1.** Summary of simple sequence repeat (SSR) and single nucleotide polymorphism (SNP) marker availability and genotyping results for linkage mapping in the GRYG, CNJ02, and CNJ04 cranberry populations following marker filtering steps and merging with previous marker datasets generated in Schlautman *et al.* (2015a) and Covarrubias-Pazaran *et al.* (2016).

| Population | # of SNPs<br>in HapMap | # SNPs<br>after<br>filtering for<br>missing<br>data<sup>a</sup> | # SNPs<br>after<br>filtering for<br>MAF<sup>b</sup> | # SNPs<br>after<br>filtering for<br>SD<sup>c</sup> | # SSRs | Total # of<br>mappable<br>markers |
| --- | --- | --- | --- | --- | --- | --- |
| GRYG | 18499<sup>d</sup> | 18374 | 8345 | 5150 | 189<sup>d</sup> | 5339 |
| CNJ02 | 15197 | 15028 | 8721 | 5564 | 629<sup>e</sup> | 6193 |
| CNJ04 | 15224 | 14818 | 8566 | 4566 | 221 | 4787 |

<sup>a</sup> SNP loci with more than 20 % missing data were excluded.
<sup>b</sup> SNP loci with a MAF less than 10 % were excluded.
<sup>c</sup> SNP loci with extreme segregation distortion were excluded.
<sup>d</sup> Generated in (Covarrubias-Pazaran *et al.* 2016)
<sup>e</sup> 541 SSRs generated in (Schlautman, Covarrubias-Pazaran, *et al.* 2015)

After using the Tassel v3.0.166 GBS analysis pipeline to filter and process the sequence reads, a total of 15197 and 15224 SNPs with good sequence tag coverage were detected that were polymorphic in the CNJ02 and CNJ04 populations, respectively; and 18499 SNPs for the GRYG population were available from the previous study (Table 1). Further filtering was performed to exclude SNP loci from each population with too much missing data (< 20 %), a minor allele frequency (MAF) less than 10 %, or extreme segregation distortion with *χ*2 p-values less than 0.00001. The 5150, 5564, and 4566 remaining SNPs were combined with the SSR marker datasets so that 5339, 6193, and 4787 markers were available for linkage mapping in the GRYG, CNJ02, and CNJ04 populations, respectively (Table 1).

### Parental Component Bin Map Construction

The number of markers mapped in each of the six parental component bin maps ranged from 1774 markers for MQ (CNJ04 maternal parent) to 2487 markers for CQ (CNJ02 paternal parent) (Table S2; Table S3; Table S4), with an average of 2080 markers per parental component map. The number of SNPs mapped in the three populations was very similar (CNJ04 = 2915, CNJ02=3326, GRYG = 3158); however, the CNJ02 population had more total markers mapped because of its larger SSR dataset (Table S5). Twelve linkage groups (LGs), corresponding to the expected haploid chromosome number in cranberry (2n=2x=24) (Hall and Galleta 1971), were retrieved for each of the cranberry parents, and total length of those twelve linkage groups varied from 845.2 cM for ST (CNJ04 maternal parent) to 1296.5 cM for MQ (CNJ02 maternal parent) (Table S2; Table S3; Table S4).

On average, the linkage groups in the six parental bin maps spanned 89.4 cM and contained 173.3 markers placed in 34.7 unique bins (Table S2; Table S3; Table S4); however, notable exceptions were present in the component maps. For example, linkage group 5 from the [BGx(BLxNL)]95 component map only contained 34 markers placed in 10 marker bins spanning 61 cM (Table S2). Population size appeared to have some effect on the average number of bins per LG, a reflection of the number of observed recombination events per LG in the population of parental gametes that fused to form the progeny, such that fewer marker bins were present in the LGs of parents from the CNJ04 population, which was approximately 1/3 and 1/5 the size of the CNJ02 and GRYG populations, respectively (Figure 2). However, there was no obvious difference in number of bins per LG in the CNJ02 and GRYG parents despite the CNJ02 population containing only half as many progeny (Figure 2).

**Figure 2.**
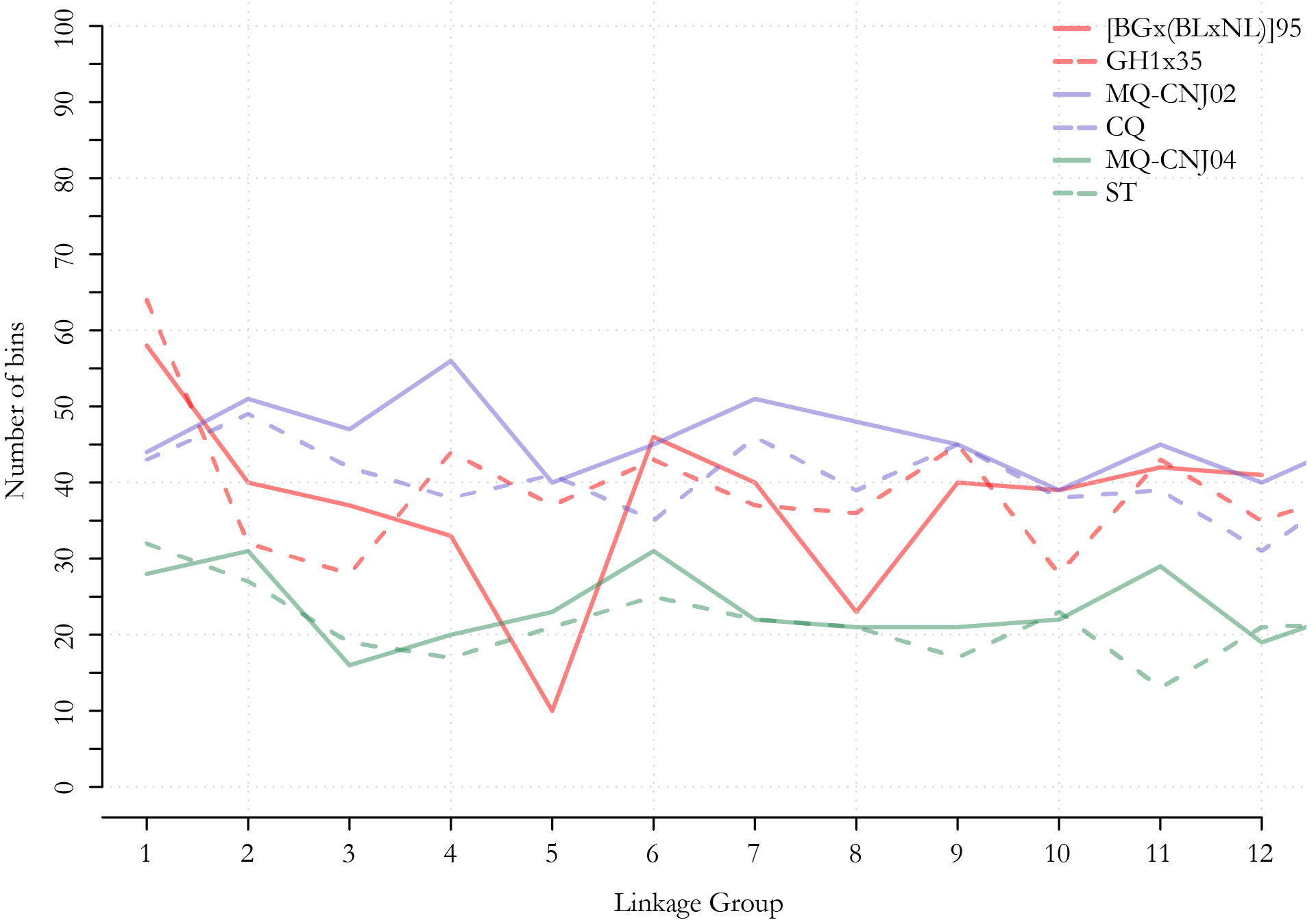
Line graphs representing the distribution of the number of recombination bins, estimated using the multiple spanning tree algorithm implemented in ASMap (Wu *et al.* 2008; Taylor and Butler 2015), in each linkage group (LG) of the six parental component bin maps constructed for three cranberry full-sib populations: GRYG (i.e. [BGx(BLxNL)]95 x GH1x35), CNJ02 (i.e. MQ x CQ), and CNJ04 (i.e. MQ x ST). The GRYG population (red) included 352 progeny; the CNJ02 population (blue) included 168 progeny, and the CNJ04 population (green) included 67 progeny.

In each parental map, the mean number of recombination events per progeny per LG ranged from 0.6 to 1.1 (Table S2; Table S3; Table S4). Comparing the number of recombination events between the 36 maternal vs. paternal LG pairs, we found that the maternal LGs had a significantly greater number of recombination events per progeny per LG (*p* < 1.0^−7^ according to a paired student t-test). The number of recombination events was greater for the maternal LG or equal to the paternal LG in 92% of the 36 pairwise comparisons, and on average 0.18 more recombination events per progeny per LG were observed in the maternal LGs. Exceptions to this trend were observed in GRYG LG 8 and CNJ04 LG 11 where there were about 0.1 more recombination events per progeny per LG in the paternal vs. maternal parent and in GRYG LG 5 where there were 0.3 more recombination events per progeny per LG in the paternal vs. maternal parent (Table S2; Table S3; Table S4).

Pair-wise Spearman rank correlations comparing marker order revealed exceptionally high levels of synteny and collinearity between the LGs in the six parental component bin maps (Table 2). An average of 57.3 markers were used in each of the 180 pair-wise comparisons of linkage groups, and a mean Spearman rank correlation of 0.993 was observed across all comparisons (Table 2). The minimum Spearman rank correlation (*r* = 0.92) was observed in comparison of LG 9 between parents [BGx(BLxNL)]95 and ST (Table 2). Most differences in marker order between linkage groups were due to markers being in a single marker bin in one parent while being in adjacent marker bins in the other parent. The high degree of synteny and collinearity among LGs in the component maps was also visually observed in alignments of the linkage maps of the parents in each of the three cranberry populations in Circos and in scatterplots of the relative position of the common markers in each map on different axes (Figure 3).

**Figure 3.**
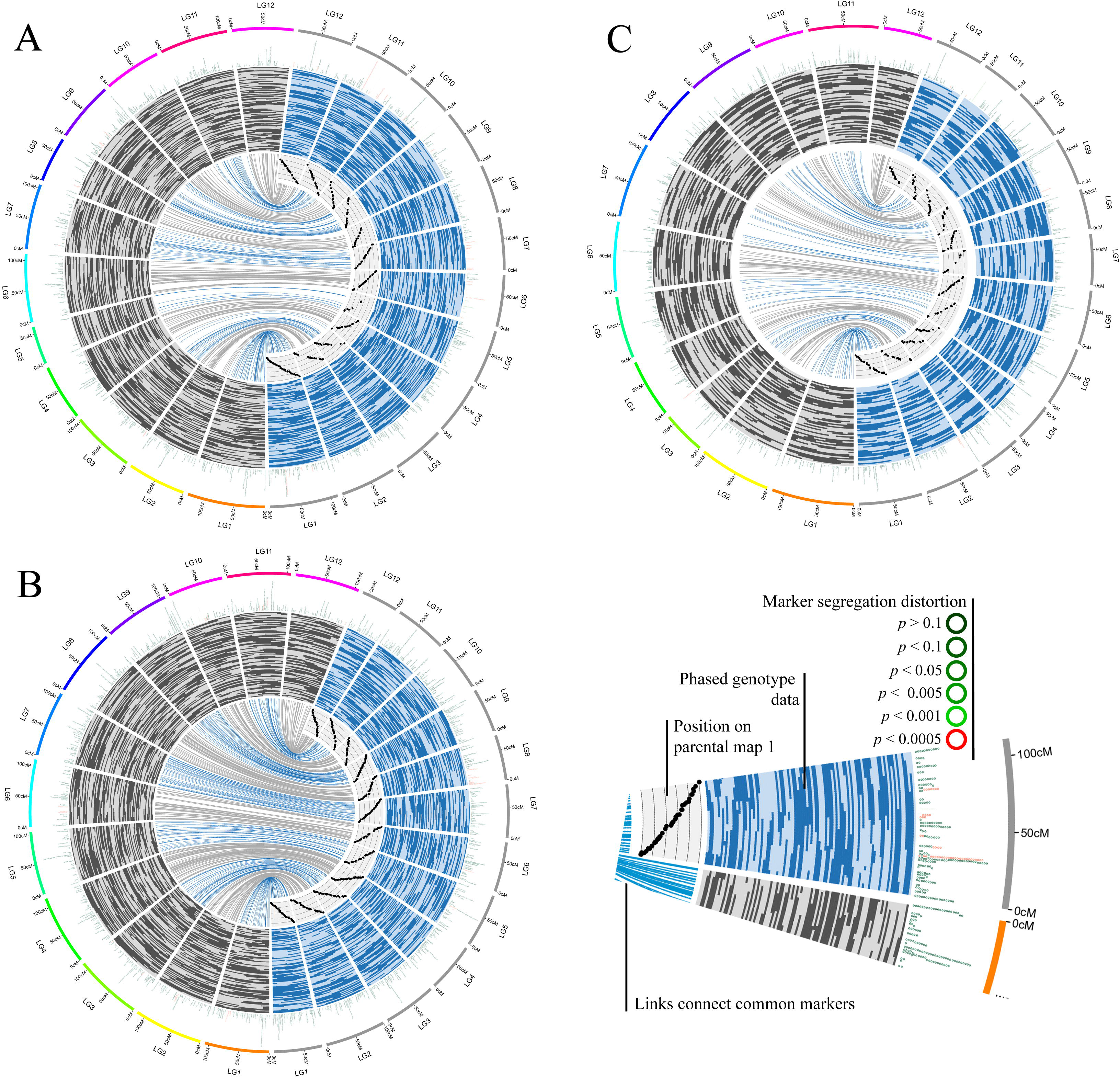
Comparisons of the LGs of the parental linkage maps for the(A) GRYG ([BGx(BLxNL)]95 x GH1x35), (B) CNJ02 (Mullica Queen x Crimson Queen), and (C) CNJ04 (Mullica Queen x Stevens) full-sib cranberry mapping populations. LGs from the maternal maps are on the left side of each circular ideogram while the paternal LGs are on the right. Links are drawn between common markers in the LGs of the two parental maps in each population. Scatterplots of the position (cM) of the common markers in the paternal map plotted on the x-axis and position in the maternal map plotted on the y-axis display collinearity of marker order. Bars display phased genotype data which shows position (cM) of phase changes (the gametic recombination) which occurred in both parents for a random subset of 60 progeny from each population. Outer ring displays the position (cM) of markers in the parental maps colored by the *χ*2 *p*-value obtained from the tests for distortion from expected Mendelian segregation ratios. Marker colors range from dark green for markers not showing distortion (*χ*2 p > 0.1) to dark red for markers showing highly significant segregation distortion (*χ*2 p < 0.0005).

**Table 2.** Pair-wise Spearman rank correlations between the linkage groups (LGs) of the six parental component maps constructed for the parents (P) of the GRYG (i.e. P1 = [BGx(BLxNL)]95 and P2 = GH1x35), CNJ02 (P3 = MQ and P4 = CQ), and CNJ04 (P5 = MQ and P6 = ST) full-sib cranberry populations. Coefficients of Coancestry (CoA), or kinship coefficients, were calculated for the parents involved in each pairwise comparison. The number of common markers available and used in each comparison is listed in parenthesis.

| LG | P1xP2 | P1xP3 | P1xP4 | P1xP5 | P1xP6 | P2xP3 | P2xP4 | P2xP5 | P2xP6 | P3xP4 | P3xP5 | P3xP6 | P4xP5 | P4xP6 | P5xP6 |
| --- | --- | --- | --- | --- | --- | --- | --- | --- | --- | --- | --- | --- | --- | --- | --- |
| 1 | 1.00<br>(91) | 1.00<br>(78) | 1.00<br>(100) | 1.00<br>(56) | 1.00<br>(75) | 1.00<br>(95) | 1.00<br>(85) | 1.00<br>(81) | 0.99<br>(71) | 1.00<br>(96) | 1.00<br>(119) | 1.00<br>(59) | 1.00<br>(52) | 0.99<br>(69) | 1.00<br>(87) |
| 2 | 1.00<br>(42) | 1.00<br>(50) | 1.00<br>(100) | 1.00<br>(46) | 1.00<br>(43) | 1.00<br>(37) | 1.00<br>(50) | 1.00<br>(29) | 1.00<br>(30) | 1.00<br>(89) | 1.00<br>(135) | 1.00<br>(32) | 1.00<br>(49) | 1.00<br>(55) | 1.00<br>(51) |
| 3 | 1.00<br>(41) | 0.99<br>(42) | 1.00<br>(56) | 0.97<br>(27) | 0.98<br>(24) | 1.00<br>(91) | 1.00<br>(37) | 0.97<br>(56) | 0.98<br>(32) | 1.00<br>(77) | 0.99<br>(97) | 0.99<br>(37) | 0.98<br>(25) | 0.99<br>(64) | 0.99<br>(44) |
| 4 | 1.00<br>(49) | 1.00<br>(66) | 1.00<br>(47) | 0.99<br>(39) | 0.99<br>(32) | 1.00<br>(68) | 1.00<br>(46) | 1.00<br>(42) | 1.00<br>(37) | 1.00<br>(96) | 1.00<br>(102) | 0.99<br>(23) | 0.99<br>(31) | 0.99<br>(62) | 0.99<br>(29) |
| 5 | 0.99<br>(15) | 0.98<br>(16) | 0.98<br>(19) | 0.95<br>(11) | 0.93<br>(8) | 1.00<br>(85) | 1.00<br>(86) | 1.00<br>(48) | 0.99<br>(34) | 1.00<br>(85) | 1.00<br>(93) | 1.00<br>(15) | 0.99<br>(35) | 1.00<br>(56) | 1.00<br>(19) |
| 6 | 1.00<br>(70) | 1.00<br>(57) | 1.00<br>(86) | 0.99<br>(43) | 1.00<br>(49) | 1.00<br>(96) | 1.00<br>(51) | 0.99<br>(83) | 0.99<br>(42) | 1.00<br>(80) | 0.99<br>(143) | 0.99<br>(49) | 1.00<br>(52) | 1.00<br>(89) | 0.99<br>(55) |
| 7 | 1.00<br>(45) | 1.00<br>(54) | 1.00<br>(50) | 0.99<br>(37) | 0.99<br>(22) | 1.00<br>(99) | 1.00<br>(44) | 0.99<br>(68) | 0.99<br>(14) | 1.00<br>(91) | 1.00<br>(122) | 0.98<br>(29) | 0.99<br>(42) | 1.00<br>(48) | 0.98<br>(34) |
| 8 | 0.99<br>(32) | 0.99<br>(29) | 0.99<br>(41) | 0.99<br>(26) | 0.99<br>(24) | 1.00<br>(35) | 1.00<br>(36) | 1.00<br>(31) | 1.00<br>(15) | 1.00<br>(83) | 1.00<br>(97) | 1.00<br>(21) | 0.99<br>(46) | 0.99<br>(55) | 1.00<br>(25) |
| 9 | 1.00<br>(84) | 1.00<br>(67) | 1.00<br>(70) | 0.99<br>(49) | 0.92<br>(54) | 1.00<br>(83) | 1.00<br>(57) | 0.99<br>(56) | 0.98<br>(37) | 1.00<br>(93) | 0.99<br>(141) | 0.97<br>(31) | 1.00<br>(39) | 0.98<br>(46) | 0.98<br>(36) |
| 10 | 1.00<br>(48) | 1.00<br>(34) | 1.00<br>(46) | 1.00<br>(31) | 0.99<br>(69) | 1.00<br>(79) | 0.98<br>(53) | 1.00<br>(59) | 0.99<br>(33) | 1.00<br>(82) | 1.00<br>(100) | 0.98<br>(27) | 0.99<br>(29) | 0.99<br>(64) | 0.99<br>(31) |
| 11 | 1.00<br>(56) | 1.00<br>(66) | 1.00<br>(77) | 0.99<br>(49) | 0.99<br>(32) | 1.00<br>(114) | 1.00<br>(69) | 0.99<br>(94) | 0.99<br>(28) | 1.00<br>(97) | 1.00<br>(159) | 0.99<br>(34) | 1.00<br>(61) | 0.99<br>(47) | 0.98<br>(43) |
| 12 | 1.00<br>(66) | 0.99<br>(48) | 1.00<br>(77) | 0.98<br>(40) | 0.98<br>(46) | 0.99<br>(85) | 1.00<br>(57) | 0.99<br>(64) | 0.97<br>(37) | 0.99<br>(77) | 0.99<br>(127) | 0.98<br>(32) | 0.99<br>(39) | 0.97<br>(75) | 0.98<br>(45) |
| CoA<sup>a</sup> | 0.094 | 0.078 | 0.094 | 0.078 | 0.063 | 0.156 | 0.031 | 0.156 | 0.063 | 0.000 | 0.500<sup>c</sup> | 0.000 | 0.000 | 0.250 | 0.000 |
| Mean | **1.00**<br>**(53.3)** | **1.00**<br>**(50.6)** | **1.00**<br>**(64.1)** | **0.99**<br>**(37.8)** | **0.98**<br>**(39.8)** | **1.00**<br>**(80.6)** | **1.00**<br>**(55.9)** | **0.99**<br>**(59.3)** | **0.99**<br>**(34.2)** | **1.00**<br>**(87.2)** | **1.00**<br>**(119.6)** | **0.99**<br>**(32.4)** | **0.99**<br>**(41.7)** | **0.99**<br>**(60.8)** | **0.99**<br>**(41.6)** |
| Total<sup>b</sup> | **639** | **607** | **769** | **454** | **478** | **967** | **671** | **711** | **410** | **1046** | **1435** | **389** | **500** | **730** | **499** |

<sup>a</sup> Coefficient of Coancestry calculated using pedigree information
<sup>b</sup> Total markers in common between the compared parental bin maps.
<sup>c</sup> self-Coancestry is 0.5 assuming no prior inbreeding

### Composite Map Construction

The high degree of collinearity between the component maps allowed for the construction of a cranberry composite map with *LPmerge* by retaining the synthetic composite map for each LG computed with the maximum interval size, *k*, which minimized the RMSE (Endelman and Plomion 2014). The composite map contained 6073 markers (636 SSRs and 5437 SNPs) spanning 1124.3 cM (Table 3; Table S6, File S2). The 12 LGs of the composite map ranged from 84.1 cM to 115.9 cM in length, and each LG contained an average of 506 (453 SNPs and 53 SSRs) markers (Table 3). The 6073 markers in the composite map corresponded to 1560 unique marker positions (bins), with an average gap of 0.7 cM between unique marker positions (Table 3). The largest gap in the composite map was on LG10 and spanned 6.6 cM; however, there were only three gaps more than 5 cM and nine gaps more than 4 cM in the entire composite map (Table S7). The composite map anchored a total of 3989 cranberry scaffolds from the Polashock *et al.* (2014) assembly totaling 21.8 Mb (4.6%) of the predicted 470 Mb cranberry genome (Table S8). Of the 3989 anchored scaffolds, 1654 contained predicted coding DNA sequences (CDS) (Table S8).

**Table 3.** Features of the cranberry composite map, constructed using the six parental component bin maps for the parents of the CNJ02, CNJ04, and GRYG populations, including the length of the linkage groups (LG), the total number of simple sequence repeat (SSR) and single nucleotide polymorphism (SNP) markers mapped, the number of unique marker positions, the number of unique marker positions containing an SSR, and the mean gap distance (cM) between unique positions.

| LG | Length<br>(cM) | # SNPs | # SSRs | Total #<br>Markers | # Unique<br>Marker<br>Positions | # Positions<br>with an<br>SSR | Mean<br>Gap<br>(cM) |
| --- | --- | --- | --- | --- | --- | --- | --- |
| LG1 | 115.9 | 588 | 50 | 638 | 158 | 37 | 0.74 |
| LG2 | 100.8 | 416 | 63 | 479 | 152 | 45 | 0.67 |
| LG3 | 92.4 | 407 | 54 | 461 | 99 | 34 | 0.94 |
| LG4 | 86.4 | 423 | 61 | 484 | 110 | 36 | 0.79 |
| LG5 | 93.1 | 405 | 35 | 440 | 110 | 24 | 0.85 |
| LG6 | 93.8 | 473 | 50 | 523 | 157 | 37 | 0.6 |
| LG7 | 97.0 | 418 | 59 | 477 | 158 | 47 | 0.62 |
| LG8 | 85.2 | 370 | 54 | 424 | 128 | 35 | 0.67 |
| LG9 | 89.9 | 532 | 54 | 586 | 112 | 33 | 0.81 |
| LG10 | 84.1 | 378 | 51 | 429 | 115 | 32 | 0.74 |
| LG11 | 95.3 | 499 | 42 | 541 | 152 | 30 | 0.63 |
| LG12 | 90.6 | 528 | 63 | 591 | 109 | 32 | 0.84 |
| **Mean** | **93.7** | **453** | **53** | **506** | **130** | **35** | **0.74** |
| **Total** | **1124.3** | **5437** | **636** | **6073** | **1560** | **422** |  |

Collinearity between the cranberry composite map and the 6 component parental bin maps was high; the mean Spearman correlation across the 72 pairwise-comparisons of marker order was *r* = 0.997 (Table 4). Collinearity between the parents of the two larger populations (i.e. CNJ02 and GRYG) and the composite map were slightly higher (i.e. *r* ≥ 0.999) than the CNJ04 parents (i.e. *r =* 0.995 and 0.991) (Table 4). Marker order variation between the composite and six component maps was highest in LG12, with perfect correlation observed only between the [BGx(BLxNL)]95 component map and the composite map (Table 4). Spearman rank correlations comparing the LGs of the composite map to the 12 LGs from the Covarrubias-Pazaran *et al.* (2016) SNP-SSR map and the Schlautman *et al.* (2015a) SSR map were also high (i.e. *r* = 0.976 and *r* = 0.996, respectively) (Table S9).

**Table 4.** Pair-wise Spearman rank correlations between the linkage groups (LGs) of the six parental component maps constructed for the parents of the GRYG ([BGx(BLxNL)]95 and GH1x35), CNJ02 (MQ and CQ), and CNJ04 (MQ and ST) full-sib cranberry populations compared to the LGs of the cranberry composite map.

| LG | [BGx(BLxNL)]95 | GH1x35 | MQ-CNJO2 | CQ | MQ-CNJO4 | ST |
| --- | --- | --- | --- | --- | --- | --- |
| LG1 | 1.00 | 1.00 | 1.00 | 1.00 | 1.00 | 1.00 |
| LG2 | 1.00 | 1.00 | 1.00 | 1.00 | 1.00 | 1.00 |
| LG3 | 1.00 | 1.00 | 1.00 | 1.00 | 0.99 | 0.99 |
| LG4 | 1.00 | 1.00 | 1.00 | 1.00 | 1.00 | 0.99 |
| LG5 | 1.00 | 1.00 | 1.00 | 1.00 | 1.00 | 0.97 |
| LG6 | 1.00 | 1.00 | 1.00 | 1.00 | 1.00 | 1.00 |
| LG7 | 1.00 | 1.00 | 1.00 | 1.00 | 1.00 | 1.00 |
| LG8 | 1.00 | 1.00 | 1.00 | 1.00 | 1.00 | 1.00 |
| LG9 | 1.00 | 1.00 | 1.00 | 1.00 | 0.99 | 0.97 |
| LG10 | 1.00 | 1.00 | 1.00 | 1.00 | 1.00 | 1.00 |
| LG11 | 1.00 | 1.00 | 1.00 | 1.00 | 1.00 | 0.99 |
| LG12 | 1.00 | 0.99 | 0.99 | 0.99 | 0.96 | 0.98 |
| **mean** | **1.000** | **0.999** | **0.999** | **0.999** | **0.995** | **0.991** |

### Genome-wide segregation distortion and centromere placement

Approximately 8 % of the markers in the six parental component bin maps displayed significant segregation distortion (SD) at the χ^2^ *p* < 0.1 level (Figure 3, Table S10). SD was not randomly distributed across the linkage groups or populations. The GH1x35 parental map contained a much higher proportion of markers displaying segregation distortion (i.e 26 %) compared to the other five parents, with as many as 81% of all markers in LG 6 displaying distortion (Figure 3, Table S10). Segregation distortion regions (SDRs) were observed in LGs of each of the six parental bin maps, and some SDRs, such as the SDR on LG 9, appeared to be present in multiple parental genomes (Figure 3, Table S10).

Phasing the genotype data and estimating RF_M_ for each marker interval from the terminal markers in both directions allowed for centromere placed on each of the 12 LGs of the component maps using the method developed in Limborg *et al.* (2016) (Table 5, Figure 4). The recombination phasing method allows for distinguishing between metacentric and acrocentric linkage groups. All cranberry linkage groups appeared to be metacentric, which is consistent with the karyotype for cranberry and diploid *Vacciniums* observed in Hall and Galleta (1971), and centromere spans averaged 18.6 cM across the 12 LGs of the six component maps (Table 5; Figure 4).

**Figure 4.**
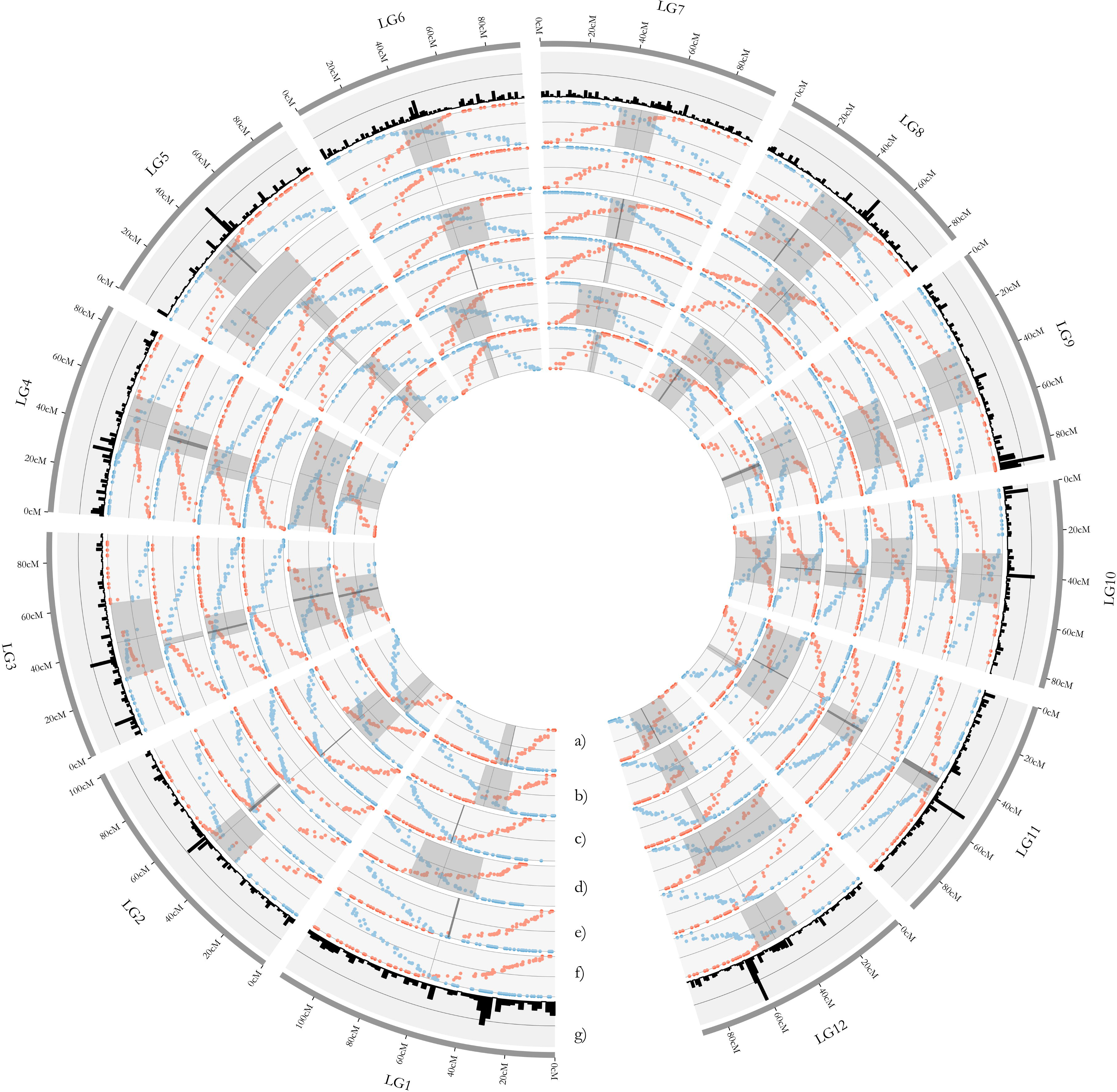
Plots of recombination frequency (RF_M_) estimated from phased genotype data by starting at the terminal markers at the beginning (red points) and end (blue points) of each linkage and recording the proportion of offspring with an observed recombination (i.e change of phase) in the interval between the terminal marker (m0) and each subsequent marker (mn) for the 12 cranberry LGs. Centromere spans (gray regions) were placed on the 12 linkage groups (LGs) of the cranberry parental component bin maps constructed for the parents of the CNJ04 population, Mullica Queen **(a)** and Stevens **(b)**; the CNJ02 population, Mullica Queen **(c)** and Crimson Queen **(d)**; and the GRYG population, [BGx(BLxNL)]95 **(e)** and GH1x35 **(f)** using the method developed in Limborg *et al.* (2016). Centromeric spans in the LGs were defined as the range (cM) extending from the intersection (dark lines) of the recombination frequency (RF_M_) estimates made from the both ends of the LG outwards until reaching the first marker with an RF_M_ = 0.45 in both directions. **(g)** Marker density in the cranberry composite map is shown to explore the relationship between marker density and centromere position.

**Table 5.** Centromere spans placed on the 12 linkage groups (LGs) of the cranberry parental component bin maps constructed for the parents of the CNJ04 population, Mullica Queen (MQ) and Stevens (ST); the CNJ02 population, MQ and Crimson Queen (CQ); and the GRYG population, [BGx(BLxNL)]95 (BGBLNL95) and GH1x35 using the method developed in Limborg et al. (2016). Centromeric spans (*x* cM – *y* cM) in the LGs were defined as the range from the intersection between the recombination frequency (RF_M_) estimates between each marker and the terminal marker on both ends of the LG extending outwards until reaching the first marker with an RF_M_ = 0.45 in both directions. The intersection point (cM) of RF_M_ estimates in each LG is provided in parentheses.

| Parent | LG |  |  |  |  |  |  |  |  |  |  |  |
| --- | --- | --- | --- | --- | --- | --- | --- | --- | --- | --- | --- | --- |
|  | 1 | 2 | 3 | 4 | 5 | 6 | 7 | 8 | 9 | 10 | 11 | 12 |
| MQ-CNJ04 | 44.3-55.0<br>(44.3) | 24.2-42.1<br>(28.2) | 20.8-58.6<br>(41.1) | 29.9-58.7<br>(36.9) | 47.1-58.5<br>(47.7) | 38.1-43.8<br>(43.5) | 42.3-49.6<br>(45.4) | 13.9-41.1<br>(23.9) | 43.8-55.3<br>(44.6) | 15.0-61.5<br>(36.6) | 46.9-53.0<br>(50.2) | 24.9-64.7<br>(49.9) |
| ST | 38.0-62.8<br>(55) | 28.2-59.2<br>(50.2) | 20.8-69.9<br>(45.5) | 4.2-79.0<br>(43.9) | 44.7-56.1<br>(48.3) | 21.0-49.8<br>(34.2) | 24.3-57.6<br>(42.3) | 20.0-62.4<br>(41.7) | 25.9-62.7<br>(38.1) | 31.7-52.8<br>(43.4) | 6.6-66.3<br>(38.1) | 35.4-60.1<br>(43.7) |
| MQ-CNJ02 | 68.8-68.8<br>(68.8) | 47.8-47.8<br>(47.8) | 42.9-42.9<br>(42.9) | 33.9-33.9<br>(33.9) | 42.4-44.7<br>(42.4) | 48.9-48.9<br>(48.9) | 42.3-45.4<br>(42.3) | 36.2-36.2<br>(36.2) | 49.3-49.3<br>(49.3) | 39.0-54.2<br>(44.0) | 38.7-38.7<br>(38.7) | 48.0-52.9<br>(48) |
| CQ | 46.1-85.2<br>(67.8) | 49.3-49.3<br>(49.3) | 33.4-47.0<br>(37.1) | 28.7-43.9<br>(33.9) | 44.7-56.1<br>(48.3) | 40.4-65.4<br>(45.9) | 38.3-53.5<br>(44.8) | 36.2-58.8<br>(49.4) | 38.1-73.2<br>(52.5) | 21.2-45.3<br>(36) | 38.1-53.6<br>(43.5) | 9.8-66.4<br>(55.3) |
| BGBLNL95 | 53.9-53.9<br>(53.9) | 50.0-52.5<br>(50.3) | 36.5-39.3<br>(36.5) | 35.6-50.1<br>(41.5) | 9.8-61.0<br>(22.4) | 44.1-44.1<br>(44.1) | 49.9-49.9<br>(49.9) | 14.7-50.2<br>(32.1) | 52.5-57.2<br>(52.5) | 37.1-44.4<br>(39.0) | 50.2-50.2<br>(50.2) | 49.6-49.6<br>(49.6) |
| GH1x35 | 58.1-58.1<br>(58.1) | 37.4-54.5<br>(50.2) | 26.0-61.0<br>(42.9) | 34.9-55.9<br>(46.7) | 26.4-46.7<br>(41.8) | 37.2-55.3<br>(45.9) | 35.7-52.9<br>(45.4) | 30.9-58.1<br>(44.7) | 31.0-56.3<br>(43.8) | 29.8-52.7<br>(39.9) | 44.6-53.6<br>(44.6) | 34.9-54.7<br>(46.3) |
| **mean** | **51.5-64.0**<br>**(58.0)** | **39.5-50.9**<br>**(46.0)** | **30-53.1**<br>**(41.0)** | **27.9-53.6**<br>**(39.4)** | **35.9-53.9**<br>**(41.8)** | **38.3-51.2**<br>**(43.7)** | **38.8-51.5**<br>**(45.0)** | **25.3-51.1**<br>**(38.0)** | **40.1-59**<br>**(46.8)** | **29.0-51.8**<br>**(39.8)** | **37.5-52.6**<br>**(44.2)** | **33.8-58.1**<br>**(48.8)** |

## DISCUSSION

Genotyping-by-sequencing is an effective strategy for simultaneous SNP discovery and genotyping in cranberry, and was used to identify thousands of polymorphic SNP loci which were transferable between inter-related full-sib mapping populations. A composite map was developed by merging the six bin maps constructed for the parents of each full-sib family, which allowed for exploration of segregation distortion regions (SDRs) and centromere placement. The map will serve as framework for future genomics studies to identify QTL regions of interest and for the continued investigation of genome organization and evolution in the genus *Vaccinium*. Furthermore, the cranberry genomic scaffolds and predicted CDS anchored in the composite map demonstrate its utility in future efforts to anchor physical maps and assist in the assembly of cranberry and *Vaccinium* genome sequences.

### Map construction

The use of three full-sibling populations of diverse genetic backgrounds facilitated an increase in the number of mapped markers (SNPs and SSRs) and resulting genome coverage in the cranberry composite map that has not been possible in past studies using single populations (Georgi *et al.* 2013; Schlautman *et al.* 2015a; Covarrubias-Pazaran *et al.* 2016). More importantly, the pedigrees of the three full-sibling populations all trace their ancestry to the seven wild cranberry selections, “The Big Seven”, made in the 19^th^ and 20^th^ centuries, which collectively account for the entire genetic base of the modern cranberry industry (Figure 1) (Eck 1990; Clark and Finn 2010; Fajardo *et al.* 2012). Therefore, the composite cranberry map affords the ideal opportunity for exploring and connecting the genetic diversity to the phenotypic diversity that has been responsible for the historical success of cranberry production. A key use for the composite map will be to allow for concurrent identification and integration of quantitative trait loci (QTL) and marker trait loci (MTL) which colocalize across populations, across environments, and across phenotypes within and among future linkage mapping, QTL mapping, and genome-wide association studies (GWAS).

The composite map will be most useful for studies within section *Oxycoccus*, and it may aid in future targeted introgression of genomic regions involved in the expression of unique metabolic pathways or disease resistance genes from the small-fruited cranberry, *V. oxycoccos*, to the American cranberry, *V. macrocarpon* (Vorsa and Polashock 2005). However, due to the high level of synteny and collinearity detected between cranberry and blueberry using cross-transferable SSRs (Schlautman 2016), information from the composite map should also be applicable to many other commercially important *Vacciniums* including blueberry (*Vaccinium* section *Cyanococcus*) and closely related sections lingonberry (*Vaccinium* section *Vitis-Idaea*) and sparkle berry (*Vaccinium* section *Batodendron*) (Lyrene 2011; Schlautman *et al.* 2016).

Two methods are commonly employed for integration of independent linkage maps. The first method uses pooled segregation and recombination information from independent maps; however, this method is computationally unfeasible for integration of high-density linkage maps containing thousands of loci (Van Ooijen 2011; Endelman and Plomion 2014; Bodénès 2015). The second method, which was used in the present study and other recent composite mapping studies in pine and oak (Chancerel *et al.* 2013; Bodénès 2015), performs map integration based on marker position rather than observed recombination such as is proposed and implemented in the *LPmerge* package (Endelman and Plomion 2014). Using *LPmerge*, the six parental component bin maps, each containing an average of 2080 markers (Table S2; Table S3; Table S4), were merged to obtain a synthetic composite map that included 6073 loci (1560 unique marker positions) covering 1124 cM on 12 LGs, making it the highest-density map in the *Vaccinium* genus and the entire Ericaceae family (Table 3, File S2). This map achieved considerably higher marker saturation (i.e. 0.7 cM mean interval between unique marker positions) than previous published cranberry linkage maps based on SSR and SNP markers (Georgi *et al.* 2013; Schlautman *et al.* 2015a; Covarrubias-Pazaran *et al.* 2016), while decreasing the total map length by 50 cM compared to the highest-density cranberry SSR map and increasing total map length by only 13 cM compared to the first cranberry SNP map (Table 3) (Schlautman *et al.* 2015a; Covarrubias-Pazaran *et al.* 2016).

The similarity in total map lengths between the current composite map and former maps reflects the robust marker-ordering and marker distance estimation achieved in construction of the six individual component maps using stringent parameters during mapping, imputing missing data and correcting potential genotyping errors, and the bin mapping strategy. Accuracy of the cranberry parental linkage maps is supported by the near perfect collinearity of the parental bin maps as quantified by Spearman rank correlations (e.g. Spearman rank correlations (*r*) exceeded 0.99 for 58 (80 %) of the 72 pair-wise comparisons of marker order in the LGs of the composite map compared to the LGs of each of the component parental bin maps and exceeded 0.95 for all 72 pair-wise comparisons), and is especially apparent when observing the recombination events as phase changes in the phased genotype data of each of the parental bin maps (Table 2, Table 4, Figure 3). Spearman rank correlations comparing the LGs of the composite map to LGs from the Covarrubias-Pazaran *et al.* (2016) SNP-SSR map and the Schlautman *et al.* (2015a) SSR map were also high (i.e. *r* = 0.976 and *r* = 0.996, respectively) and provided further evidence of the composite map quality (Table S9).

Previous high-density linkage mapping studies in outcrossing populations have often excluded the biparental markers (i.e. hk x hk segregation) because of difficulties they cause during mapping and because of assumptions that they are less informative for linkage mapping (Ward *et al.* 2013; Bodénès 2015); however, including the biparental SNPs in this study greatly increased the number of total markers mapped and the number of markers in common between component maps. All genetic maps (i.e. composite and component maps) in this study could be directly compared because of the high frequency of transferability of SNP markers across parents and populations. A total, 2921 (54%) of the SNPs in the composite map were mapped in two or more populations and 1040 (19%) of the SNPs were mapped in all three populations (Table S5). The number of markers in common between linkage groups in the parental maps appeared to be correlated (*r* = 0.74) with the degree of shared ancestry between the parents. For example, there were, on average, 80.1 markers in common per LG for parents with a coefficient of coancestry greater than 0.1, while parents with a coefficient of coancestry less than 0.1 had an average of 49 markers in common per LG (Table 2, Figure 1). A high degree of synteny and collinearity was visually observed in alignments of the parental maps in each population (Figure 3), and the mean Spearman rank correlation of *r* = 0.993 was remarkably high in comparisons of marker order, based on an average of 57.3 markers per LG, across all 180 pairwise comparisons of LGs in the parental component maps (Table 2). The observed transferability of SNP markers across the five cranberry parents from three populations suggests that GBS will be an effective marker discovery and genotyping platform for exploring cranberry genetic diversity and population structure, and aided by the placement of more than 5000 SNPs in the composite map, will be useful in estimating linkage disequilibrium (LD) in cranberry diversity panels for genome-wide association studies (GWAS).

In addition to the more than 5000 SNP markers mined from GBS data and mapped in this study, the composite map also included 636 SSRs, which represents the largest collection of mapped SSR markers available for cranberry and an important genetic resource for future cranberry breeding efforts (Table 3; Table S1). Of the 1540 unique marker positions in the composite map, 27% include one or more SSR markers (Table 3). Within the Mullica Queen and Crimson Queen parental bin maps for the CNJ02 population, an average of 53% of the unique marker bins include an SSR marker suggesting that a larger number of marker positions in the composite map could have included one or more SSR markers if more SSR data been available for the other two full-sib populations (Table S3). The large number of positioned SSRs, and their apparent distribution throughout the entire cranberry genome, should allow them to be useful in marker-assisted seedling selection of marker trait loci (MTL) in cranberry using multiplexing panels like those developed in (Schlautman 2016). Despite the obvious benefits of simultaneous marker discovery and genotyping afforded by GBS, the high cost per sample does not yet justify its use for large-scale genotyping of thousands of seedlings for selection of a few MTL in a specialty crop such as cranberry. A better way to allocate resources will be to utilize the SSRs, which are likely to be polymorphic in much more diverse backgrounds than SNPs (Hamblin *et al.* 2007), to select for must-have MTL in large populations for simple traits, and then use genotyping-by-sequencing to further explore the genetic diversity or perform genomic selection in the MTL selected individuals (Dekkers 2007; Collard *et al.* 2008; Singh and Singh 2015).

### Recombination rate variation

Comparisons across the six parental linkage maps developed in this study highlighted interesting differences that are related to the rates of recombination in each parent and may have important implications for cranberry breeding and genetic studies. First, the average number of marker bins per linkage group, which reflects the number of unique recombination events in the parental gametes, was lower in the parents of the CNJ04 population (23 bins per parent per LG), than in the GRYG and CNJ02 populations (38 bins and 43.5 bins per parent per LG, respectively) (Table S2; Table S3; Table S4; Figure 2). The fewer bins observed in the parents of the CNJ04 population was expected and is likely due to its small size (67 progeny), suggesting that breeders may need less genetic data (DNA markers) to capture all the recombination history in small biparental breeding populations. Unexpectedly, there were similar numbers of marker bins per LG in the CNJ02 and GRYG populations although the GRYG population (352 progeny) was much larger than the CNJ02 (168 progeny) population. This observation indicates that cranberry genotypes can exhibit different rates of recombination or different numbers of recombination hot and cold spots. Differential rates of recombination frequencies between cranberry genotypes were also observed in (Schlautman *et al.* 2015a; Covarrubias-Pazaran *et al.* 2016), and both genotypic and environmental effects on genetic recombination were observed in tetrad analysis of cranberry reciprocal translocation heterozygotes (Ortiz and Vorsa 2004). Finally, the smaller number of bins observed in the CNJ04 population suggests that increasing cranberry breeding population size from ~70 progeny to ~150 progeny can increase the number of unique recombination events observed; however, the similar number of bins observed in the large (GRYG) and medium size (CNJ02) populations suggests that cranberry breeders could better allocate their resources by planting many medium size populations (~150 progeny) rather than a few large populations (~350 progeny) because they will not necessarily detect significantly more unique recombination events in the larger populations.

The second interesting observation is that the paternal linkage groups were consistently shorter than the maternal linkage groups, and there were fewer phase changes (i.e. recombination events) per progeny per LG in the paternal gametes than in the maternal gametes (Table S2; Table S3; Table S4, Figure 3). Sex-specific differences in recombination rates have previously been observed in cranberry (Schlautman *et al.* 2015a; Covarrubias-Pazaran *et al.* 2016), blueberry (Schlautman 2016), apple (Maliepaard *et al.* 1998), and olive (Sadok *et al.* 2013). It is uncertain whether these observed differences in recombination rates between sexes are more than coincidental differences in patterns and rates of recombination in the five cranberry parents. However, we speculate that the differential recombination rates in male and female gametes observed in *Vaccinium* linkage mapping studies may be a true consequence resulting from a combination of common practices employed by cranberry and blueberry breeders during population development, differences between microsporogenesis and megasporogenesis, and the unique pollen morphology of the *Vaccinium* genus.

During megasporogenesis, three of the four megaspores disintegrate, and only one megaspore survives; conversely, all four microspores survive during microsporogenesis (Fehr 1991). Previous studies in *Vaccinium* species including cranberry revealed that pollen, rather than being shed as single grains which is common in flowering plants, are shed as groups of four pollen grains (i.e. the four microspores) called tetrads which are derived from the same meiotic division (Eck 1990; Roper and Vorsa 1997). This has important consequences considering that, during meiosis, chiasma almost always occur between only two of the four chromatids on each side of the centromere (Roeder 1997). Therefore, assuming *Vaccinium* chromosomes are metacentric (Hall and Galleta 1971), for any single chromosome in a *Vaccinium* pollen tetrad (i.e. a total of eight chromosome arms), four of the eight chromosome arms in the four haploid gametes (i.e. four microspores) represent meiotic recombinants, but in reality, there are only two unique positions of recombination because the reflection of each recombinant chromosome arm exists as one of the three remaining recombinant chromosome arms in the pollen tetrad.

In addition, pollen germination studies in both cranberry and blueberry have revealed that all four pollen grains are viable in the majority of *Vaccinium* pollen tetrads (Huang and Johnson 1996; Cane 2009), and that fruit set does not increase after loading more than 8 tetrads on a cranberry stigma which contain an average of 32 ± 4 ovules (Sarracino and Vorsa 1991; Cane and Schiffhauer 2003). Consequently, it is highly likely that any seed within a *Vaccinium* fruit shares a paternal meiotic history with one or more seeds in the same fruit; however, because of the disintegration of three of the four megaspores, no seed in a fruit shares a maternal meiotic history with any other seed in the same fruit. The result of this phenomena (high *Vaccinium* microspore fertility with pollen shed as tetrads), combined with the fact that *Vaccinium* breeders and geneticists generally only make a few crosses (i.e. harvest seeds from a few fruits) to generate breeding and/or linkage mapping populations, may explain the reduced rate of recombination observed in the paternal versus maternal gametes in this study. Specifically, in a population of 40 cranberry full-sib progeny with a meiotic history tracing to 10 cranberry pollen tetrads (i.e. 40 haploid paternal gametes), assuming strong cross-over interference, a maximum of 20 unique recombinant chromosome arms out of 80 could be observed for any single paternal metacentric chromosome in the full-sib progeny; conversely, a maximum of 80 out of 80 unique recombinant chromosome arms could be observed for the same maternal metacentric chromosome in the full-sib progeny. Although future studies will be needed to confirm this hypothesis, these results suggest that *Vaccinium* breeders and geneticists should consider harvesting seeds from as many crosses as possible (i.e. more fruits) during population development to ensure equal representation of recombination in both the maternal and paternal genomes. Although pollen tetrad formation could be a challenge in *Vaccinium* breeding and genetics; pollen tetrad analysis, which not feasible in most higher Eukaryotes, has been highly advantageous and used extensively in genetic studies of *Arabidopsis* mutants to detect every genetic change between chromatids, to simplify genetic map construction, to define centromere positions, and to quantify crossover and chromatid interference (Preuss *et al.* 1994; Copenhaver *et al.* 2002; Brieuc *et al.* 2014). Furthermore, it has already been useful in detecting chromosomal rearrangements, specifically reciprocal translocations, in cranberry cultivars (Ortiz and Vorsa 2004).

### Genome characterization

SNP and SSR loci present within nuclear genome scaffolds from the Polashock *et al.* (2014) assembly were used anchor 3989 nuclear scaffolds containing 21.8 Mb (4.6%) of the predicted 470 Mb cranberry genome in the composite map (Table S8). Although the present composite map anchored 1500 more nuclear scaffolds than the previous SNP linkage map, those 1500 scaffolds only represented a 1.9% increase in the total Mb of cranberry genome anchored (Covarrubias-Pazaran *et al.* 2016). This is reflective of the sheer number of scaffolds (i.e. 229,745 scaffolds) and their size (i.e. N50 = 4,237 bp); and suggests the need for a higher quality cranberry genome assembly (Polashock *et al.* 2014).

Segregation distortion affects accuracy of linkage map construction by introducing errors in map distance estimation and marker order, and thus, could affect mapping quantitative trait loci (QTL) (Zhang *et al.* 2010). Deviations from expected Mendelian inheritance were widespread throughout the cranberry genome, and ~8 % of the markers positioned in the parental component maps displayed significant segregation distortion (i.e. *p* < 0.1) according to chisquare tests (χ^2^) with one degree of freedom (Figure 3, Table S10). Previous high-density linkage mapping studies have sometimes observed that distorted markers are not always randomly distributed, but rather, are grouped together in segregation distortion regions (SDRs) (Yin *et al.* 2004; Wang *et al.* 2012a; Bodénès 2015; Covarrubias-Pazaran *et al.* 2016). Likewise, apparent SDRs were observed in the present study in all six of the parental component maps (Figure 3, Table S10). Many of the SDRs are unique to each parental cranberry genome, as was previously observed in oak and palm (Ting *et al.* 2014; Bodénès 2015). However, there is apparent overlapping between some SDRs in the six parents (e.g. SDR in LG9 for the [BGx(BLxNL)]95 and Stevens parents, and SDR in LG8 for the [BGx(BLxNL)]95 and Crimson Queen parents), which could represent biologically significant phenomena such as gametic competition, gametophytic selection, or sterility (Wang *et al.* 2005; Bloom and Holland 2012; Xu *et al.* 2013). The identification of SDR regions in this study provides a framework to further study the cause of this phenomena in cranberry and to improve experimental designs for association mapping studies and marker-assisted selection strategies.

There have been minimal efforts to characterize chromosomal structures, such as centromeres, in cranberry, and modern cytogenetic approaches such as fluorescent in situ hybridization (FISH) have not yet been attempted. Centromeres are central components of chromosome architecture that play fundamental roles in the regulation and crossover formation during meiosis (Roeder 1997; Zickler and Kleckner 1999); and therefore, knowledge of centromere location is critically important in attempts to manage the occurrence of meiotic recombination in plant breeding (Wijnker and de Jong 2008). However, many high-density linkage mapping studies in cranberry and other commercially important crops have not attempted or failed to place centromeres onto the linkage groups (Wang *et al.* 2012b; Castro *et al.* 2013; Ting *et al.* 2014; Bodénès 2015; Covarrubias-Pazaran *et al.* 2016), which could limit their future applicability in interpretations of genome organization, genome divergence, and of the genomic architecture of adaptive or economically important traits (Limborg *et al.* 2016). Therefore, we utilized the recombination phasing method outlined in Limborg *et al.* (2016) and effectively applied in Mckinney *et al.* (2016) to define centromere regions in the cranberry linkage groups (Table 5; Figure 4).

In general, the Limborg *et al.* (2016) recombination phasing method appeared to work well in cranberry. Centromeres were placed on all twelve cranberry linkage groups in each of the six parental component maps (Table 5; Figure 4). All linkage groups were identified as metacentric, consistent with a previous cranberry karyotype presented in Hall and Galleta (1971). The centromere regions in the parental linkage groups, although sometimes large (i.e. an average of 18.6 cM across LGs), were similar in size to those observed in Mckinney *et al.* (2016). The centromere regions appeared to have the same general location across linkage groups in the parental maps, providing further confidence in the estimation of their positions. Interestingly, observed marker densities were often higher in centromere regions and had reduced recombination rates (i.e. recombination “coldspots”) compared to the rest of the respective linkage groups (Figure 4). The knowledge of centromere location generated herein should facilitate future meiotic studies in cranberry exploring crossover interference and recombination “hotspots” and “coldspots”, and could potentially be useful for developing cranberry crop improvement strategies for managing meiotic recombination and for overcoming barriers to recombination in cranberry chromosomes (Martinez-Perez and Moore 2008).

In conclusion, genotyping-by-sequencing has been shown to be a highly efficient means for SNP marker discovery and genotyping in cranberry. A large proportion of the genotyping-by-sequencing based SNP loci were polymorphic and transferable between three full-sib cranberry populations, which allowed for the construction of the first a high-density composite linkage map in cranberry composed of both SNP and SSR markers. The stringent parameters used during component map construction, and the remarkable collinearity observed between the six component maps and the composite map suggests that estimation of marker position and distance was performed in an accurate, reproducible manner. The large number of cross-transferable mapped markers allowed for characterization of cranberry genomic architecture such as the detection of centromeric regions, and we foresee that the composite map and the marker data will be used extensively for future QTL detection, genome-wide association studies, and development of molecular-assisted breeding strategies in cranberry.

## AUTHOR CONTRIBUTIONS

BS, GCP, NV, JP, and JZ conceived the study. JZ, MI, NV, JP supervised the research. BS and GCP prepared samples and conducted DNA genotyping. BS and GCP formatted genotypic data, performed imputations, and constructed the linkage and composite maps. BS performed all statistical analyses. BS and LDG identified putative centromere regions. EG, NV, JP, JJC, and MI provided intellectual contributions and/or germplasm resources. BS and JZ wrote the manuscript. BS and LDG prepared the final figures. The manuscript was read, edited, and approved by all authors.

## Acknowledgements and Funding

JZ and BS wish to express their gratitude through 1 Cor 10:31. This project was supported by USDA-SRCI under Grant 2008-51180-04878; USDA-NIFA-AFRI Competitive Grant USDA-NIFA-2013-67013-21107: USDA-ARS (project no. 3655-21220-001-00 provided to JZ); WI‐ DATCP (SCBG Project #14-002); National Science Foundation (DBI-1228280); Ocean Spray Cranberries, Inc.; NJ Cranberry and Blueberry Research Council; Wisconsin Cranberry Growers Association; Cranberry Institute. BS was supported by the Frank B. Koller Cranberry Fellowship Fund for Graduate Students; GCP and LDG were supported by the Consejo Nacional de Ciencia and Tecnologia (CONACYT, Mexico). MI was supported by the USDA National Institute of Food and Agriculture, Hatch project 1008691.

---

**File S1.** Parameters used in the Tassel v3.0.166 reference-based genotyping-by-sequencing (GBS) pipeline for processing raw sequence data and calling SNPs in the resulting sequence data generated for the parents and progeny of the CNJ02, CNJ04, and GRYG cranberry populations.

**File S2.** Position of single nucleotide polymorphism (SNP) and simple sequence repeat (SSR) markers in the cranberry composite and six parental maps and the associated cranberry genomic scaffold sequence from Polashock *et al.* (2014).

**File S3.** SNP Genotype data for the six parental component bin maps ([BGx(BLxNL)]95, GH1x35, Mullica Queen (CNJ02), Crimson Queen, Mullica Queen (CNJ04), and Stevens).

**Table S1.** Simple sequence repeat (SSR) markers used to genotype the parents and progeny of the CNJ02, CNJ04, and GRYG full-sib cranberry populations in this study, their primer sequences, their publication of origin, their NCBI ID, and their position (cM) in the linkage groups (LG) of the cranberry composite map constructed herein.

**Table S2.** Features of the parental bin maps and linkage groups (LGs) constructed for the maternal parent (M), [BGx(BLxNL)]95, and the paternal parent (P), GH1x35, for the GRYG full-sib mapping population using simple sequence repeats (SSRs) and single nucleotide polymorphisms (SNPs).

**Table S3.** Features of the parental bin maps and linkage groups (LGs) constructed for the maternal parent (M), Mullica Queen, and the paternal parent (P), Crimson Queen, for the CNJ02 full-sib mapping population using simple sequence repeats (SSRs) and single nucleotide polymorphisms (SNPs).

**Table S4.** Features of the parental bin maps and linkage groups (LGs) constructed for the maternal parent (M), Mullica Queen, and the paternal parent (P), Stevens, for the CNJ04 full-sib mapping population using simple sequence repeats (SSRs) and single nucleotide polymorphisms (SNPs).

**Table S5.** Total number of markers, by linkage group (LG), mapped in the parental component bin maps for each of the three cranberry full-sib populations (i.e. GRYG, CNJ02, and CNJ04).

**Table S6.** Statistic generate during composite map construction with the six parental component bin maps from the cranberry GRYG, CNJ02, and CNJ04 populations using LPmerge (Endelman and Plomion 2014). The max interval size, *k*, which minimized the root mean square error (RMSE) is displayed along with the corresponding RMSE and standard deviation (SD).

**Table S7.** The number of gaps between unique marker positions in the cranberry consensus map exceeding 1, 2, 3, 4, and 5 cM in length by linkage group (LG).

**Table S8.** Total number of cranberry scaffold, by linkage group (LG), from the Polashock *et al.* (2014) assembly containing SNPs or SSRs that were anchored in the cranberry composite map. The number of anchored scaffolds containing predicted coding DNA sequences (CDS) and total number of base pairs (bp) contained within those scaffolds is also provided.

**Table S9.** Pair-wise Spearman rank correlations between the linkage groups (LGs) the cranberry composite map and previous cranberry linkage maps (Schlautman *et al.* 2015; Covarrubias-Pazaran *et al.* 2016).

**Table S10.** Proportion of markers in the linkage groups (LGs) of the 6 parental component maps (i.e. [BGx(BLxNL)]95, GH1x35, Mullica Queen (MQ), Crimson Queen (CQ), and Stevens (ST) from the GRYG, CNJ02, and CNJ04 populations) that display significant segregation distortion from the expected Mendelian genotype rations according to *χ*2 tests at the *p* < 0.1 level.

